# Structural Connectivity Between the Locus Coeruleus and Nucleus Basalis of Meynert Relates to Dual-Task Performance

**DOI:** 10.64898/2026.07.20.739186

**Authors:** AmirHussein Abdolalizadeh, Yan Deng, Karsten Witt, Christiane M. Thiel

**Affiliations:** Biological Psychology Lab, Department of Psychology, School of Medicine and Health Sciences, Carl von Ossietzky University Oldenburg, Oldenburg, Germany; Department of Neurology, School of Medicine and Health Science, Carl von Ossietzky University Oldenburg, Oldenburg, Germany; Research Center Neurosensory Science, Carl von Ossietzky University Oldenburg, Oldenburg, Germany

**Keywords:** Locus coeruleus, Basal nucleus of Meynert, fMRI, Diffusion MRI, Dual-task

## Abstract

The noradrenergic locus coeruleus (LC) and the cholinergic nucleus basalis of Meynert (nBM) are key hubs of ascending neuromodulatory systems that shape large-scale brain dynamics. However, the behavioral relevance of the structural and functional connectivity between these nuclei remains poorly understood. Here, we investigated whether LC-nBM structural or functional connectivity is related to cognitive-motor dual-task performance in healthy younger and older adults. Fifty-four participants (36 older, 18 younger) underwent diffusion MRI, resting-state fMRI, and behavioral assessment using an MRI-compatible cognitive-motor dual-task paradigm. LC-nBM structural connectivity was estimated using tractography, whereas resting-state functional connectivity was quantified as Fisher z-transformed correlations between LC and nBM time series. LC-nBM structural connectivity was better explained by a quadratic rather than a linear or cubic age model, indicating non-linear age-related variation, whereas functional connectivity showed no significant age-related association. Higher LC-nBM structural connectivity was associated with greater cognitive dual-task cost, but not with motor dual-task cost or single- or dual-task reaction times. This association was not moderated by age group and was not statistically explained by attentional and executive performance as measured by the Test of Attentional Performance. These findings suggest that LC-nBM structural connectivity is selectively associated with cognitive-motor interference, potentially reflecting a neuromodulatory pathway that constrains the balance between task-specific stabilization and flexible cross-domain coordination during a cognitive-motor dual-tasking.

## 1. Introduction

Cognitive-motor dual-tasking is a common feature of everyday behavior, requiring simultaneous coordination of cognitive control and motor execution. Walking while monitoring the environment, responding to cues, or making decisions places demand on attentional allocation, response selection, and the flexible coordination between higher cognitive and sensorimotor systems. Performance decline in such conditions, commonly referred to as dual-task costs, is therefore considered a sensitive marker of cognitive-motor interference (Leone et al. 2017). In older adults and clinical populations, increased dual-task costs have been linked to neurodegenerative decline (Bishnoi and Hernandez 2021; Yang et al. 2020).

At the neural level, successful dual-task performance relies on the ability to transiently integrate distributed cognitive and motor systems while preserving task-specific processing. This maps onto the broader principle that adaptive behavior requires a dynamic balance between network integration and segregation, with integration supporting coordination across systems and segregation preserving specialized processing within task-relevant networks (Shine et al. 2016). Although not studied in a dual-task context, previous work suggests that successful motor performance is associated with more segregated network configurations, whereas successful cognitive performance is associated with more integrated network states (Cohen and D’Esposito 2016). Cognitive-motor dual-tasking may therefore place competing demands on large-scale brain organization, requiring the simultaneous coordination of processes that benefit from partially different network regimes. Dual-task costs can therefore be understood not only in terms of limited resources, central bottlenecks, or executive task coordination demands (Leone et al. 2017), but also as a behavioral expression of how effectively the brain regulates the integration-segregation balance.

Ascending neuromodulatory systems play an important role in regulating brain states that support various cognitive functions by influencing distributed cortical and subcortical networks and shaping neural dynamics (Avery and Krichmar 2017). Importantly, several of these systems, including the noradrenergic Locus Coeruleus (LC) and cholinergic nucleus basalis of Meynert (nBM), are known to undergo age-related changes and have been implicated in cognitive decline and neurodegenerative disease (Shafiee et al. 2024; Liu et al. 2019). They are particularly relevant for adaptive control, influencing the integration-segregation balance through their widespread cortical and subcortical projections (Shine 2019), thereby regulating adaptive gain, task engagement, and attentional selection (Aston-Jones and Cohen 2005; Sarter et al. 2005). Clinical studies have also supported the role of neuromodulatory systems in dual-task performance. Observational studies in patients with Parkinson’s disease have reported that lower nBM gray matter density is associated with poorer gait during a cognitive-motor dual task (Nishida et al. 2024; Dalrymple et al. 2021). This view is complemented by interventional studies using cholinergic medications, which have shown improved gait features during a cognitive-motor dual-task in Alzheimer’s disease (Shimura et al. 2021) and Parkinson’s disease (Henderson et al. 2016). In contrast to nBM, evidence from direct LC neuroimaging linking the noradrenergic neuromodulatory system to cognitive-motor dual-task performance is limited. However, pupil dilation, an indirect marker of LC activity (Murphy et al. 2014), increases with cognitive control demands and postural/dual-task balance difficulty (Kahya et al. 2018; 2021). Together, these findings indicate that, at the behavioral level, cognitive-motor dual-tasking can be influenced separately by cholinergic and, potentially, noradrenergic neuromodulatory pathways.

Although the LC and nBM are often studied separately, anatomical and physiological evidence suggest that they form an interacting noradrenergic-cholinergic circuit (Shine 2019). Experimental rodent tracing studies have demonstrated LC projections to basal forebrain structures, predominantly ipsilateral but partially bilateral (España and Berridge 2006; Jones et al. 1977); moreover, LC projections to the nBM appear to excite basal forebrain cholinergic neurons, while direct reciprocal projections from the nBM to the LC have not been demonstrated (Zaborszky and Gombkoto 2017; Schwarz and Luo 2015; Záborszky et al. 1993). This architecture suggests that LC activity may influence large-scale brain dynamics both directly and through its connections to the basal forebrain cholinergic nuclei. Supporting this view, Munn et al. (2021) showed that phasic activity in LC and nBM is temporally linked to large-scale changes in cortical network topology and brain-state energy landscapes, with LC-related activity associated with greater network integration and landscape flattening, and nBM activity associated with more segregated dynamics. Extending this framework, Taylor et al. (2022) used diffusion MRI tractography in healthy young adults and showed that greater LC-nBM structural connectivity was associated with higher network-level integration irrespective of phasic LC or nBM activity, as well as flattening of attractor landscape dynamics. These findings suggest that LC-nBM connectivity may provide a conduit for regulating transitions between integrated and segregated brain states. Given that cognitive-motor dual-tasking places substantial demands on the dynamic coordination of cognitive and motor systems, and because these functions are known to decline with age, it provides a particularly relevant framework for investigating the role of LC-nBM connectivity in healthy aging.

In the present study, we combined diffusion MRI tractography and resting-state fMRI to quantify LC-nBM structural and functional connectivity in healthy older and younger adults. We first examined whether LC-nBM connectivity showed age-related linear or nonlinear trajectories. Furthermore, we tested whether LC-nBM connectivity was associated with cognitive and motor dual-task costs derived from an MRI-compatible cognitive-motor dual-task paradigm. Finally, given that dual-task interference may depend on attentional and executive control, we examined whether performance on neuropsychological measures indexing these functions statistically mediated the association between LC-nBM connectivity and dual-task cost. We hypothesized that LC-nBM connectivity would be related to dual-task cost, reflecting the role of noradrenergic-cholinergic circuitry in supporting cognitive-motor control under competing task demands.

## 2. Methods

### 2.1. Participants

A total of 61 healthy participants were recruited for this study through local advertisements. Participants were sampled to form two age groups: older adults aged 50-80 years (n = 41; 19 males; mean age (SD) = 67.17 (7.54)) and younger adults aged 20-40 years (n = 20; 11 males; mean age (SD) = 28.0 (4.87)). Inclusion criteria were age within the predefined range for each group, right-handedness, and the ability to walk independently without assistive devices. Participants were required to report good physical and mental health, assessed using the Short Form-12 Health Survey (SF-12; Ware et al. 1996). Exclusion criteria included self-reported neurological, psychiatric, or motor disorders. Participants taking centrally acting medication or having standard MRI contraindications were excluded. We further excluded six participants due to the lack of either single- or dual-task reaction times, as their responses inside the scanner were not recorded. An additional subject was excluded due to failed tractography, resulting in a final sample of 54 participants (36 older and 18 younger adults).

The study protocol was approved by the research ethics committee of the Carl von Ossietzky University of Oldenburg and conducted in accordance with the Declaration of Helsinki (Drs.EK/2020/062-02). Written informed consent was obtained from all participants.

### 2.2. Behavioral Measurements

#### 2.2.1. Cognitive-Motor Dual-Task Performance

Behavioral indices of cognitive-motor dual-task performance were obtained using an MRI-compatible pedaling paradigm. While task-related fMRI activity associated with this paradigm has been reported previously (Deng et al. 2026), the present study examined whether individual differences in dual-task performance were associated with LC-nBM structural and resting-state functional connectivity. Briefly, participants completed a cognitive Go/NoGo task, a motor pedaling task, and a combined dual-task condition during a task fMRI acquisition. Cognitive and motor dual-task costs were calculated as 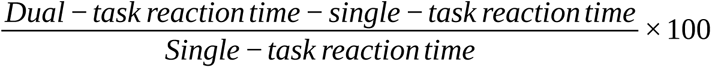.

#### 2.2.2. Test of Attentional Performance (TAP-M)

Following MRI acquisitions, the participants performed the Test of Attentional Performance - Mobility (TAP-M, version 1.3.2; Zimmermann and Fimm 2004), a computerized battery assessing attentional and executive functioning. Based on their relevance to dual-task performance, we included measures of distractor suppression, executive control, divided attention (auditory and visual), response inhibition, and Go/NoGo in subsequent analyses.

### 2.3. MRI Acquisition

All imaging data were acquired on a 3T Siemens Prisma scanner at the Carl von Ossietzky University of Oldenburg using a 64-channel head coil. A structural T1-weighted magnetization-prepared rapid gradient echo (MPRAGE) image was acquired (TR/TE/TI = 2000/2.07/952 ms, flip angle = 9°, voxel size = 0.75 mm isotropic). For structural connectivity, a diffusion-weighted spin-echo echo-planar image (EPI) was acquired (TR/TE = 3247/79 ms, phase encoding direction = AP, bandwidth = 1775 Hz/pixel, partial Fourier = 6/8, multiband acceleration factor = 3, voxel size = 1.5 mm isotropic). A set of interspersed shells with b-values of 300, 700, and 2000 s/mm^2^ was acquired in 8, 32, and 64 diffusion-encoding directions. 13 interspersed non-diffusion-weighted b0 images were also acquired. Additionally, we acquired a b0 image with the opposite phase-encoding direction (PA) for later susceptibility distortion correction. For functional connectivity, a resting-state T2*-weighted EPI was acquired (TR/TE = 850/30 ms, multiband acceleration factor = 4, partial Fourier = 8/8, and voxel size = 2.52 × 2.52 × 2.00 mm). For each subject, 560 volumes were acquired, totaling 8 minutes of scan time. During the resting-state fMRI scan, participants were instructed to keep their eyes open and fixate on a white circle presented on a black background at the center of the screen.

To correct for susceptibility distortions, a B0 field map was acquired and included two magnitude images and a phase-difference image (TR/TE1/TE2 = 400/5.19/7.65 ms, flip angle = 60°, matrix size = 64 × 64, slice thickness = 3.0 mm, and inter-slice spacing = 0.99 mm).

### 2.4. Structural MRI Analysis

We performed T1 image analysis and surface reconstruction using FastSurfer v2.3.0 (https://github.com/Deep-MI/FastSurfer; Henschel et al., 2020; 2022) to obtain relevant anatomical segmentations and cortical surface reconstructions for subsequent diffusion and resting-state analyses. Additionally, we utilized FreeSurfer’s hippocampal and thalamic nuclei segmentation modules on FastSurfer’s outputs (Iglesias et al. 2015; 2018), which were later used to improve tractography.

### 2.5. Diffusion MRI Analysis

#### 2.5.1. Diffusion MRI preprocessing

Diffusion MRI (dMRI) was preprocessed using QSIprep version 0.22.0 (RRID: SCR_002502; Cieslak et al. 2021). A detailed explanation of the dMRI preprocessing is provided in the supplementary materials. Briefly, the data were denoised, then subjected to Gibbs ringing artifact removal and bias field correction. Then, motion, susceptibility distortion, and eddy-current correction were applied. Finally, the dMRI data were resampled to AC-PC-aligned T1 space, generating upsampled, preprocessed dMRI data in AC-PC space with 1.25 mm isotropic voxels. For three participants without a reverse-phase-encoding b0 image, Synb0-DisCo was used to synthesize a distortion-free b0 image from the T1-weighted image, which was then integrated in the subject’s BIDS directory to support susceptibility-distortion correction of the diffusion MRI data (Schilling et al. 2019).

#### 2.5.2. Regions of Interest

We utilized the binarized LC meta-mask (https://osf.io/sf2ky/; Dahl et al. 2022), which was created by aggregating data from several published LC maps and provides a robust mask suitable for independent studies. Using the warps available in the OSF repository and each subject’s QSIprep output, we transformed right and left LC masks to MNI152NLin2009cAsym space, then into subject AC-PC space in a single step using *antsApplyTransform* (Avants et al. 2008).

For nBM, we thresholded and binarized the probabilistic cytoarchitectonic map at 40% (Zaborszky et al., 2021; 2008). This threshold was selected based on prior work investigating inconsistencies across atlas-based nBM masks, in which a 40% threshold yielded a relatively spatially consistent region of interest across atlas implementations while preserving anatomical coverage for tractography (Wang et al. 2022). As nBM masks were already in MNI152NLin2009cAsym space, they were transformed to subject AC-PC space using warps generated in QSIprep.

#### 2.5.3. Whole-Brain Tractography

We used MRtrix version 3.0.4 (RRID: SCR_006971; Tournier et al. 2019) for further analysis of the preprocessed dMRI data. To minimize the extra-axonal influence on fiber orientation estimates, we limited our analysis to the shell with a b-value of 2000 s/mm^2^ (Genc et al. 2020). Unsupervised 3-tissue response functions were estimated using the *Dhollander* approach (Dhollander et al. 2016). After calculating a study-average response function for each tissue type, a single-shell 3-tissue constrained spherical deconvolution method was utilized to obtain fiber orientation distributions (FOD) using MRtrix3Tissue (https://3Tissue.github.io, Dhollander and Connelly 2016), a fork of MRtrix3. FODs were further normalized by applying log-domain intensity normalization (Dhollander et al. 2021).

A 5-tissue-type (5TT) image was then generated using the Hybrid Surface and Volume Segmentation (HSVS) algorithm (Smith et al., 2020) from the FastSurfer-analyzed structural MRI data. Given that structural MRI analysis was performed in the original T1 space, an additional transformation of the 5TT image to the AC-PC space was applied (Avants et al. 2008). Finally, a whole-brain anatomically-constrained probabilistic tractography with the 2nd-order integration over FOD algorithm (iFOD2) was done to generate 50 million streamlines with the following arguments: Step-size = 0.625 mm, min-max length = 5 - 300 mm, FOD cutoff = 0.06, dynamic seeding, and backtrack enabled (Tournier J.-D., Calamante F, Connelly A. 2010; Smith et al. 2012; 2015).

#### 2.5.4. Quantitative LC-nBM Tractography

The quantitative LC-nBM tractography procedure was adapted from Taylor et al. (2022), whose work provided the main structural connectivity framework that motivated the present study. Following their approach, targeted LC-nBM streamlines were generated, merged with a whole-brain tractogram, and weighted using spherical-deconvolution-informed filtering of tractograms (SIFT2; Smith et al. 2015) to derive a quantitative and more biologically grounded structural connectivity estimate. We introduced minor modifications to improve reproducibility and comparability within our sample, including hemisphere-specific ipsilateral tractography and ensuring inclusion of more direct ipsilateral LC-nBM connections prior to SIFT2 weighting.

The AC-PC 5TT image was edited to include LC masks as subcortical and nBM masks as cortical gray matter. Using the edited 5TT image, two additional *tckgen* calls were used to generate 15,000 streamlines for each hemisphere, providing the hemisphere’s LC mask as the seed region and the ipsilateral nBM as the target region. Ipsilateral targeting was chosen a priori based on experimental tract-tracing data demonstrating that LC projections to basal forebrain structures are predominantly ipsilateral (∼80-85%), while retaining a smaller contralateral component (España and Berridge 2006; Jones et al. 1977). Other tractography parameters were kept the same as whole-brain tractography, except that the maximum length was set to 100 mm and *the seed_unidirectional* flag was enabled. Generated tracts were upsampled by a factor of 3, and the median fiber lengths of the right and left LC-nBM tracts for each subject were extracted using *tckstats*. Then, we used *tckedit* to extract 7450 from LC-nBM tracts for each hemisphere, using the median length as the maximum fiber length. We chose 7450 instead of the median value of 7500 because one subject had fewer than 7500 fibers at the median length. This specific step ensured the inclusion of a constant number of fibers for all participants connecting the LC and nBM, and selected fibers that almost directly connect the LC to the nBM, while avoiding erroneous streamlines (Figure 1).

**Figure 1.**
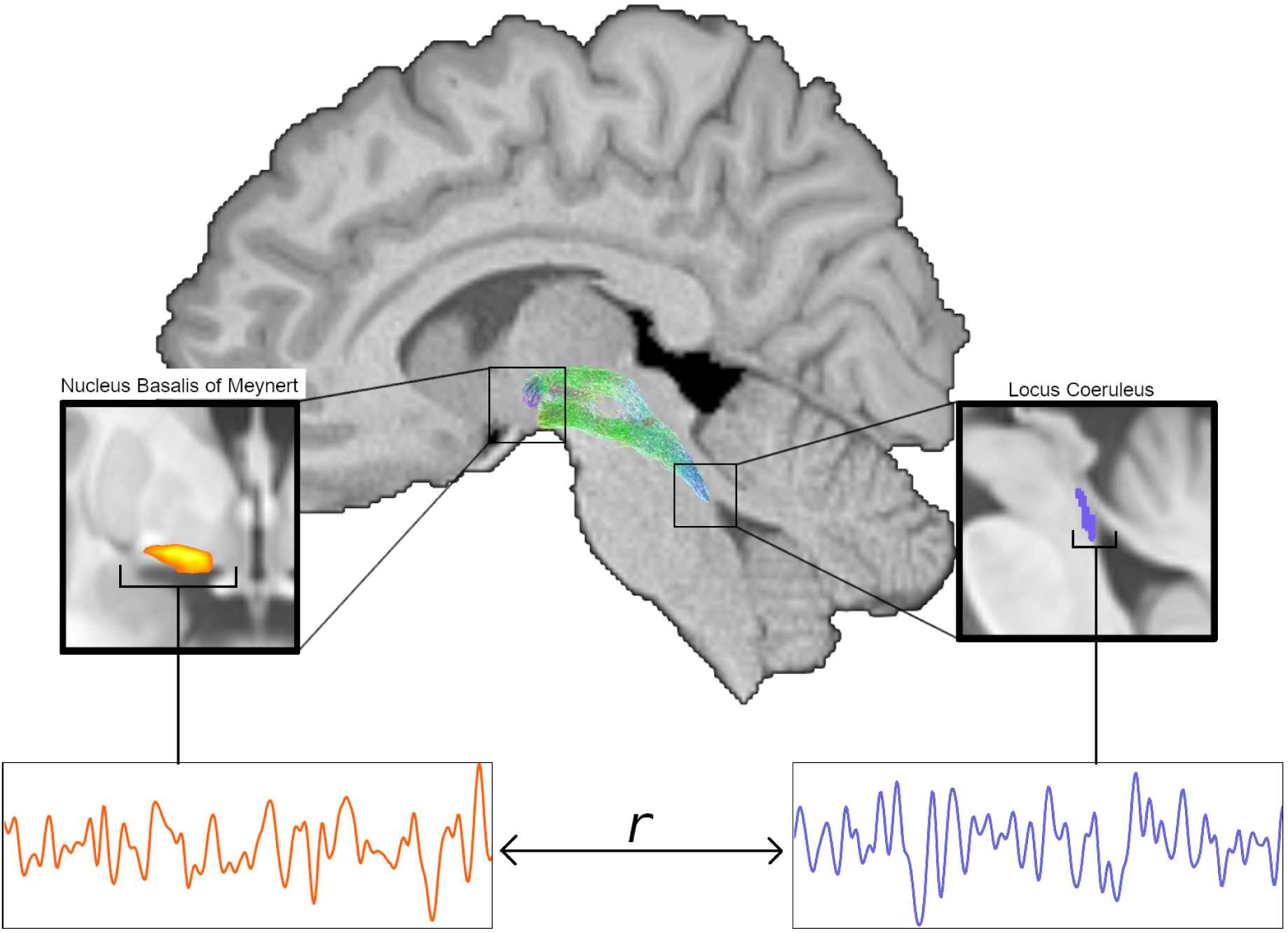
The structural and functional connectivity between locus coeruleus (LC) and nucleus basalis of Meynert (nBM). The streamlines connecting LC (right) to nBM (left) for a single subject are shown at the top. Fiber orientation is color-coded: green for dorsal-ventral and blue for cranial-caudal. Average BOLD time series were extracted from LC and nBM and used to calculate functional connectivity (bottom).

The LC-nBM tracts were then merged with the 50,000,000 whole-brain tractography streamlines, yielding a single tractogram of 50,014,900 streamlines. To enhance the biological accuracy of the tractography-based connectivity estimates, we applied SIFT2 (Smith et al. 2015) to the merged tractogram. This method assigns a weight to each streamline, improving the quantitative interpretation of the connectome (Calamante 2019; Smith et al. 2022). LC-nBM tracts for each hemisphere were again extracted from the merged tractogram using the same criteria as in the *tckedit* step above, with SIFT2-derived weights. The SIFT2 weights for each LC-nBM tract were summed to yield a single LC-nBM structural connectivity measure. One subject was excluded due to failure of the left LC-nBM tractography.

### 2.6. Functional MRI Analysis

#### 2.6.1. Resting-state functional MRI preprocessing and postprocessing

Resting-state functional MRI (rsfMRI) data were preprocessed using fMRIprep version 25.0.0 (RRID: SCR_016216; Esteban et al. 2019; Markiewicz et al. 2025) and postprocessed by XCP-D version 0.10.7 (Mehta et al. 2024) to generate a preprocessed and denoised rsfMRI in MNI152NLin2009cAsym space. The boilerplates and detailed explanation of the steps are provided in the supplementary material.

In brief, raw rsfMRI data were head-motion, slice-timing, and susceptibility distortion corrected, followed by coregistration to the subject’s T1w brain. Standard confound estimates, including framewise displacement (FD), DVARS, global signals, motion parameters, and anatomical CompCor components, were derived during preprocessing. Three participants lacked fieldmap images. We used SynBOLD-DisCo, which estimates an undistorted BOLD reference from each participant’s T1-weighted anatomical image (Yu et al. 2023). These images were then integrated into the BIDS directories of these three subjects for use during the distortion-correction step. In the postprocessing step, volumes with FD > 0.5 mm were flagged for censoring. For comparability, we followed the nuisance regression strategy used by (Jacobs et al. 2018): the top five aCompCor components from white matter and cerebrospinal fluid, as well as six motion parameters and their derivatives. Data were band-pass filtered (0.008-0.1 Hz), denoised, and censored to minimize motion- and physiology-related artifacts.

#### 2.6.2. LC-nBM Functional Connectivity

LC masks were transformed to MNI152NLin2009cAsym space using the transformation matrix and warp provided in the mask’s OSF repository. Given that nBM masks were already provided in the target standard space, no additional transformation was performed. Bilateral LC masks were combined into a single LC ROI, and bilateral nBM masks were combined into a single nBM ROI. Mean BOLD time series were extracted from each ROI, and LC-nBM functional connectivity was quantified as the Pearson correlation between the two time series, followed by Fisher’s r-to-z transformation (Figure 1).

### 2.7. Statistical Analysis

Statistical analysis was performed using R version 4.5.1 (R Core Team 2025). Demographic and behavioral differences between younger and older adults were assessed using the *compareGroups* package (Subirana et al. 2014), with tests automatically selected based on variable type. To characterize age-related trajectories of LC-nBM connectivity, associations between age and the structural and functional connectivity measures were examined using linear, quadratic, and cubic regression models. Model comparisons were performed using ANOVA to determine whether higher-order terms significantly improved model fit.

Associations between LC-nBM connectivity and cognitive-motor dual-task performance were tested using robust linear regression (*lmrob*) from the *robustbase* package (Maechler et al. 2024) to reduce the influence of outliers and heteroscedasticity. Separate models were fitted for cognitive and motor dual-task costs as outcome variables, with LC-nBM structural or functional connectivity entered as the predictor of interest, while adjusting for age and sex.

Mediation analysis was performed as a follow-up to the robust regression analysis to evaluate whether significant associations between LC-nBM connectivity and dual-task cost were statistically explained by behavioral measures of attentional and executive performance as indexed by TAP-M tests. Median reaction times of distractibility, executive control, divided attention (auditory and visual), and correct Go responses were then tested separately as a mediator of the significant association identified in the previous step, with the relevant LC-nBM connectivity measure entered as the predictor and the corresponding dual-task cost measure entered as the outcome. Mediation analysis was performed using the *mediation* package (Tingley et al. 2014), with both the mediator and outcome models adjusted for age and sex. Indirect effects were estimated using nonparametric bootstrapping with 10000 simulations. The average causal mediation effect (ACME), average direct effect (ADE), total effect, and proportion mediated were reported with 95% confidence intervals and corresponding *p*-values.

## 3. Results

### 3.1. Sample and Behavioral Characteristics

Older adults exhibited slower cognitive single-task performance. Under dual-task conditions, they showed reduced cognitive dual-task costs but increased motor dual-task costs relative to younger adults (Table 1). Older adults also showed slower performance on three of the selected neuropsychological tests, namely distractibility, auditory divided attention, and Go/NoGo. In contrast, no significant group differences were observed in either structural or functional LC-nBM connectivity measures. Sex distribution was comparable across age groups.

**Table 1.**
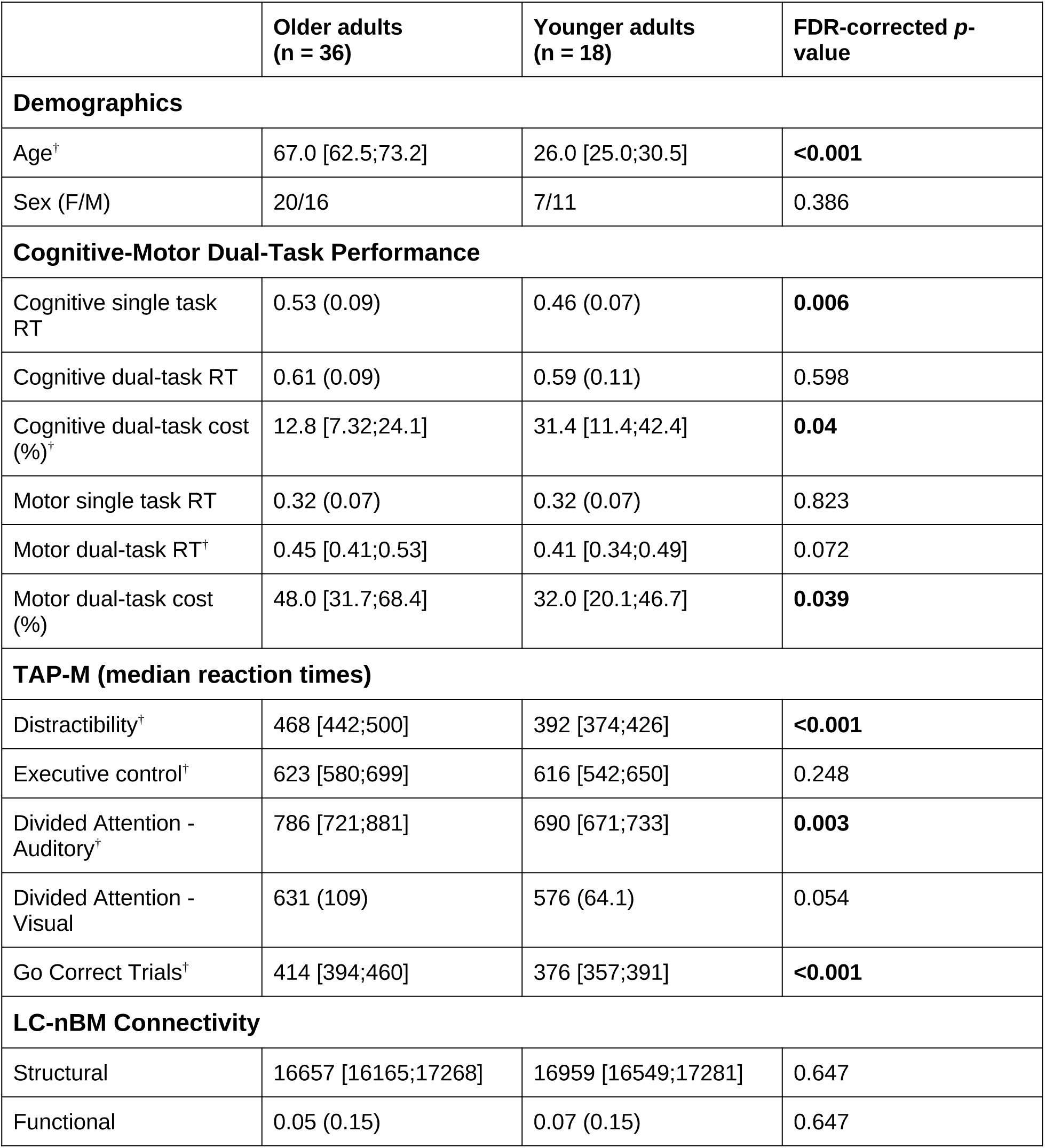
Group comparison results between older and younger adults. The variables are represented as mean (SD). The variables marked by ^†^ are represented by their median [interquartile ranges]. RT = Reaction Time, LC = Locus Coeruleus, nBM = nucleus basalis of Meynert.

|  | Older adults<br>(n = 36) | Younger adults<br>(n = 18) | FDR-corrected *p*-<br>value |
| --- | --- | --- | --- |
| **Demographics** |  |  |  |
| Age<sup>†</sup> | 67.0 [62.5;73.2] | 26.0 [25.0;30.5] | **<0.001** |
| Sex (F/M) | 20/16 | 7/11 | 0.386 |
| **Cognitive-Motor Dual-Task Performance** |  |  |  |
| Cognitive single task RT | 0.53 (0.09) | 0.46 (0.07) | **0.006** |
| Cognitive dual-task RT | 0.61 (0.09) | 0.59 (0.11) | 0.598 |
| Cognitive dual-task cost (%)<sup>†</sup> | 12.8 [7.32;24.1] | 31.4 [11.4;42.4] | **0.04** |
| Motor single task RT | 0.32 (0.07) | 0.32 (0.07) | 0.823 |
| Motor dual-task RT<sup>†</sup> | 0.45 [0.41;0.53] | 0.41 [0.34;0.49] | 0.072 |
| Motor dual-task cost (%) | 48.0 [31.7;68.4] | 32.0 [20.1;46.7] | **0.039** |
| **TAP-M (median reaction times)** |  |  |  |
| Distractibility<sup>†</sup> | 468 [442;500] | 392 [374;426] | **<0.001** |
| Executive control<sup>†</sup> | 623 [580;699] | 616 [542;650] | 0.248 |
| Divided Attention - Auditory<sup>†</sup> | 786 [721;881] | 690 [671;733] | **0.003** |
| Divided Attention - Visual | 631 (109) | 576 (64.1) | 0.054 |
| Go Correct Trials<sup>†</sup> | 414 [394;460] | 376 [357;391] | **<0.001** |
| **LC-nBM Connectivity** |  |  |  |
| Structural | 16657 [16165;17268] | 16959 [16549;17281] | 0.647 |
| Functional | 0.05 (0.15) | 0.07 (0.15) | 0.647 |

### 3.2. Nonlinear age-related trajectories of LC-nBM connectivity

Although LC-nBM connectivity did not differ significantly between younger and older adults, age-related effects may not be adequately captured by a simple group comparison. We therefore examined whether structural and functional LC-nBM connectivity showed linear or nonlinear associations with age across the full sample.

For the LC-nBM structural connectivity, no significant linear association with age was observed (β = −1.55, SE = 6.85, *p* = 0.822, adjusted R^2^ = −0.018). However, adding the quadratic age term significantly improved model fit (ΔR^2^ = 0.132, *F* = 7.76, *p* = 0.007). In the quadratic model, the age^2^ term was significant (β = 1.45, SE = 0.52, *p* = 0.007, adjusted R^2^ = 0.099). Adding a cubic term did not further improve model fit (ΔR^2^ = 0.009, *p* = 0.476). This suggests a nonlinear, quadratic relationship between age and LC-nBM structural connectivity (Figure 2A).

**Figure 2.**
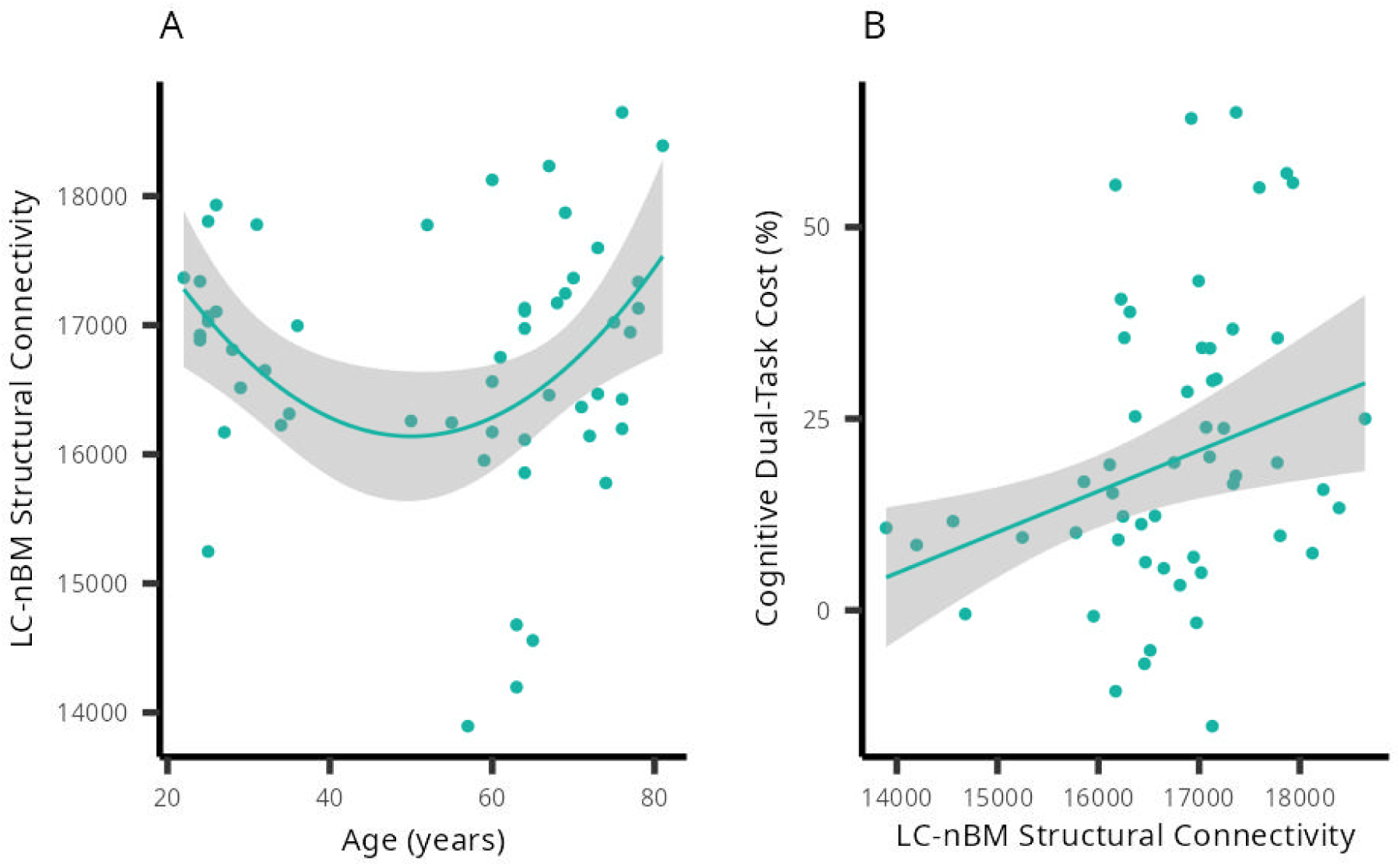
LC-nBM structural connectivity: age-related trajectories and association with cognitive dual-task cost. (**A**) LC-nBM structural connectivity shows a quadratic association with age, and (**B**) it is positively associated with cognitive dual-task cost.

For LC-nBM functional connectivity, no significant age-related association was observed. The linear model was non-significant (β = 0.0003, SE = 0.0010, *p* = 0.768), and adding quadratic or cubic terms did not significantly improve model fit (both *p* > 0.29).

### 3.3. LC-nBM Connectivity and Dual-Task Costs

To address our next research question, we examined whether individual differences in LC-nBM structural and functional connectivity were associated with cognitive and motor dual-task costs. Robust linear regression models were fitted across the full sample, with cognitive and motor dual-task costs as dependent variables and connectivity measures as predictors of interest, while adjusting for age and sex.

Of the four tested connectivity-behavior associations, only LC-nBM structural connectivity was significantly associated with dual-task performance. Specifically, higher structural connectivity was associated with greater cognitive dual-task cost (β = 0.005, SE = 0.0018, *p* = 0.006, adjusted R^2^ = 0.137; Figure 2B). No significant associations were observed for resting-state functional connectivity or for motor dual-task cost outcomes (all *p* > 0.05).

To examine the specificity of this association, we additionally tested whether LC-nBM connectivity was associated with single-and dual-task reaction times in the cognitive and motor conditions. No significant associations were observed for either structural or functional connectivity measures across these outcomes (all *p* > 0.05). Together, these findings indicate that structural, but not functional, LC-nBM connectivity is selectively associated with the additional cognitive demands of performing the cognitive and motor tasks simultaneously, rather than with general performance in either task alone.

To determine whether age group moderated the association between LC-nBM structural connectivity and cognitive dual-task cost, an additional robust linear model was fitted, including the interaction term *connectivity × group*, adjusting for sex. The interaction was not significant (β = 0.0026, SE = 0.008, *p* = 0.759), indicating that the connectivity-behavior relationship did not differ between older and younger adults. Across groups, higher LC-nBM structural connectivity remained significantly associated with greater cognitive dual-task cost (β = 0.004, SE = 0.0016, *p* = 0.01, adjusted R^2^ = 0.127), suggesting that this association is similar across age groups.

### 3.4. TAP-M as a mediator

Given that dual-task costs have been linked to attentional and cognitive control processes (Leone et al. 2017), we performed a series of mediation analyses to test whether the association between LC-nBM structural connectivity and cognitive dual-task cost was statistically explained by neuropsychological measures indexing distractibility, executive control, divided attention, and response inhibition. None of these measures showed a significant indirect effect (all ACME *p*-values > 0.12; Supplementary Table 1). In all models, the direct effect of LC-nBM structural connectivity on cognitive dual-task cost remained significant (all ADE *p*-values [FDR-corrected] < 0.05), indicating that the observed connectivity-behavior relationship was not statistically mediated by any of the assessed attentional or executive function measures.

## 4. Discussion

This study investigated the behavioral relevance of connectivity between the noradrenergic LC and cholinergic nBM for cognitive-motor dual-task performance in healthy aging. We found that higher LC-nBM structural connectivity was associated with greater cognitive dual-task cost, whereas functional LC-nBM connectivity was not associated with dual-task costs. Notably, this association was specific to the cognitive component of dual-task cost, remained stable across younger and older adults, and was not explained by attentional or executive function measures.

Given the well-established age-related vulnerability of both the noradrenergic LC and the cholinergic basal forebrain systems (Shafiee et al. 2024; Liu et al. 2019), the absence of a simple age-related decline in LC-nBM structural connectivity was unexpected. LC-nBM structural connectivity did not differ significantly between younger and older adults but showed a nonlinear, U-shaped relationship with age. Nonlinear age-related trajectories of LC-nBM functional connectivity have been reported previously (Jacobs et al. 2018). In addition, nonlinear age-related changes in diffusion MRI measures of white matter integrity have been observed across major fiber pathways (Behler et al. 2021; Davis et al. 2009), suggesting that age-related changes in structural connectivity are often not adequately captured by simple linear models.

In contrast to the structural findings, we did not observe significant age-related trajectories in LC-nBM functional connectivity. This differs from previous reports of nonlinear age-related changes in LC-nBM functional resting-state connectivity (Jacobs et al. 2018). One possible explanation is the different age range sampled across studies, as Jacobs et al. included a middle-aged group that may have been critical for capturing nonlinear age-related changes. Methodological differences in MRI acquisition and LC localization may have further contributed to the observed differences in findings. In addition, structural and functional connectivity capture different aspects of the noradrenergic-cholinergic system. Whereas tractography-derived measures reflect relatively stable anatomical pathways, resting-state functional connectivity is more strongly influenced by transient fluctuations in neural activity and brain state dynamics (Park and Friston 2013).

Although our age-related findings were observed in structural rather than functional LC-nBM connectivity, they align with the broader idea that interactions between these neuromodulatory systems may be relevant to age-related changes in cognitive control. For example, Cicero et al. (2025) reported age-related differences in task-related LC-nBM functional connectivity during an auditory Go/NoGo task, with stronger LC-nBM coupling during NoGo trials in younger adults but during Go trials in older adults, suggesting that LC-nBM coupling may vary with both age and task demands.

A growing body of work has highlighted the functional importance of both the noradrenergic LC and the cholinergic basal forebrain systems for adaptive behavior. The noradrenergic LC system regulates adaptive gain and behavioral flexibility through its widespread cortical and subcortical projections (Aston-Jones and Cohen 2005). In contrast, cholinergic signaling from the basal forebrain has been linked to enhanced selective processing and increased signal-to-noise ratios in task-relevant neural representations (Sarter et al. 2005). More recently, these neuromodulatory influences have been considered within a network neuroscience framework, in which interactions between the LC and nBM have been proposed to regulate transitions between integrated and segregated network states, thereby supporting flexible adaptation to changing task demands (Shine 2019).

For example, Munn et al. (2021) showed that phasic activity in the human LC and nBM was associated with distinct patterns of large-scale network reconfiguration, suggesting complementary roles in regulating transitions between brain states. Importantly, Taylor et al. (2022) further demonstrated that individual differences in LC-nBM structural connectivity are associated with the magnitude of these network reconfigurations. Our finding that LC-nBM structural connectivity was associated with cognitive dual-task cost, therefore, extends prior work by linking this pathway to behavioral performance. Cognitive-motor dual-tasking is particularly relevant in this context because successful motor and cognitive performance, in a single-task paradigm, have been associated with different large-scale network configurations, with motor performance relying more strongly on segregated network states and cognitive performance on integrated network states (Cohen and D’Esposito 2016; Shine et al. 2016).

Unexpectedly, however, higher LC-nBM structural connectivity was associated with greater rather than lower cognitive dual-task costs. Based on previous work linking LC-nBM connectivity to neuromodulator-driven network reconfiguration (Taylor et al. 2022), we initially expected stronger connectivity to facilitate the flexible coordination of cognitive and motor systems, thereby reducing dual-task interference, given its association with greater large-scale network integration and a flatter brain-state energy landscape. Although Taylor et al. demonstrated that LC-nBM structural connectivity is associated with these large-scale network dynamics, the behavioral significance of this relationship remained unresolved.

Given our findings, it is possible that the ability to induce large-scale network reconfigurations is not advantageous in all behavioral contexts. A flatter energy landscape may facilitate transitions between brain states, and these transitions may involve shifts between more integrated and more segregated network configurations. During cognitive-motor dual-tasking, this may be particularly relevant. Within the framework proposed by Cohen and D’Esposito (2016), successful cognitive performance has been associated with more integrated network states, whereas successful motor performance has been linked to more segregated network configurations. Cognitive-motor dual-tasking may therefore place competing demands on large-scale brain networks by requiring the simultaneous coordination of processes that benefit from different network regimes. From this perspective, stronger LC-nBM connectivity may not necessarily facilitate performance. Instead, under certain circumstances, such as cognitive-motor dual-tasking, stronger coupling between these neuromodulatory systems could make it more difficult to maintain the stabilization required for successful cognitive task performance amid concurrent motor demands.

Although speculative, this interpretation is consistent with the observation that LC-nBM connectivity was specifically related to cognitive dual-task cost, not to performance during the corresponding motor or cognitive single-task conditions. Importantly, this association was also not statistically mediated by attentional and executive control measures derived via the TAP-M battery. This suggests that LC-nBM connectivity may not primarily reflect general attentional capacity or cognitive flexibility, but rather mechanisms that become relevant when cognitive and motor systems must be coordinated simultaneously.

Despite the use of multimodal imaging, behavioral assessment, and a dual-task setup, several limitations remain. First, although our interpretation is based on the current models of neuromodulator-driven network dynamics, we did not directly measure task-related network integration or segregation. Future studies that combine LC-nBM structural connectivity with task-based measures of large-scale network organization will therefore be important for more directly testing the proposed mechanism. Second, although LC-nBM structural connectivity showed a nonlinear association with age, our sample comprised two age groups rather than a continuous lifespan cohort. Replication in larger lifespan samples will therefore be required to better characterize age-related trajectories.

## 5. Conclusions

This study provides evidence that LC-nBM structural connectivity is relevant for cognitive-motor interference in healthy adults. Unexpectedly, higher LC-nBM structural connectivity was associated with greater cognitive dual-task cost, suggesting that stronger structural coupling between noradrenergic and cholinergic neuromodulatory systems does not necessarily confer an advantage during cognitive-motor dual-tasking. Together, these findings extend previous work on LC-nBM-related brain-state dynamics by demonstrating the behavioral relevance of this pathway.

## 6. Data Availability Statement

The raw imaging data will be made available via the Oldenburg Research Data Repository (DARE) upon publication. The scripts used for statistical and neuroimaging analyses of the data are available at https://github.com/amirhusseinab/LC_nbm_dualtask.

## 7. CRediT authorship contribution statement

**Conceptualization:** AA, CT; **Data curation:** YD, AA; **Formal analysis:** AA; **Funding acquisition:** CT, KW; **Investigation:** AA, CT; **Methodology:** AA, CT, YD, KW; **Project administration:** AA, CT; **Resources:** CT; **Software:** AA; **Supervision:** CT; **Visualization:** AA, **Writing - original draft:** AA, **Writing - review and editing:** CT, YD, KW.

## 10. Acknowledgments

The analysis was performed at the high-performance cluster ROSA, located at the University of Oldenburg (Germany) and funded by the DFG through its Major Research Instrumentation Programme (INST 184/225-1 FUGG) and the Ministry of Science and Culture (MWK) of the Lower Saxony State.

