## Supplementary materials for "Structural Connectivity Between the Locus Coeruleus and Nucleus Basalis of Meynert Relates to Dual-Task Performance"

**Detailed neuroimaging analysis.** The boilerplates automatically generated by the neuroimaging analysis tools used in the current manuscript: QSIPrep, fMRIPrep, and XCP-D.
**Supplementary Table 1.** Mediation analyses testing whether individual TAP-M reaction time measures mediate the association between LC-nBM structural connectivity and cognitive dual-task cost.
**References.**

### Supplementary methods

#### Diffusion MRI Preprocessing

Preprocessing was performed using QSIPrep 0.22.2, which is based on Nipype 1.8.6 (RRID:SCR_002502; Esteban et al., 2025; Gorgolewski et al., 2011).

The anatomical reference image was reoriented into AC-PC alignment via a 6-DOF transform extracted from a full Affine registration to the MNI152NLin2009cAsym template. A full nonlinear registration to the template from AC-PC space was estimated via symmetric nonlinear registration (SyN) using antsRegistration (Avants et al., 2008). Brain extraction was performed on the T1w image using SynthStrip (Hoopes et al., 2022) and automated segmentation was performed (Tournier et al., 2019) using SynthSeg (Billot et al., 2023) from FreeSurfer version 7.3.1.

Any diffusion-weighted images with a b-value less than 100 s/mm$^2$ were treated as a b=0 image. MP-PCA denoising as implemented in MRtrix3's dwidenoise (Veraart et al., 2016) was applied with an auto-voxel window. After MP-PCA, Gibbs unringing was performed using MRtrix3's mrdegibbs (Kellner et al., 2016). Following unringing, the mean intensity of the DWI series was adjusted so that all the mean intensities of the b0 images matched across each separate dMRI scanning sequence. B1 field inhomogeneity was corrected using dwibiascorrect from MRtrix3 with the N4 algorithm (Tustison et al., 2010) after corrected images were resampled.

FSL's eddy was used for head motion correction and Eddy current correction (Andersson and Sotiropoulos, 2016). Eddy was configured with a q-space smoothing factor of 10, a total of 5 iterations, and 1000 voxels used to estimate hyperparameters. A quadratic first-level model and a linear second-level model were used to characterize Eddy current-related spatial distortion. Q-space coordinates were forcefully assigned to shells. Field offset was attempted to be separated from subject movement. Shells were aligned post-eddy. Eddy's outlier replacement was run (Andersson et al., 2016). Data were grouped by slice, only including values from slices determined to contain at least 250 intracerebral voxels. Groups deviating by more than 4 standard deviations from the prediction had their data replaced with imputed values. Slice-to-volume correction was estimated with temporal order 6, 5 iterations, trilinear interpolation, and lambda=1.000 (Andersson et al., 2017).

Data was collected with reversed phase-encode blips, resulting in pairs of images with distortions going in opposite directions. FSL's TOPUP (Andersson et al., 2003) was used to estimate a susceptibility-induced off-resonance field based on b0 reference images with reversed phase encoding directions. The TOPUP-estimated fieldmap was incorporated into the Eddy current and head motion correction interpolation. Dynamic susceptibility distortion correction was applied with 10 iterations, lambda=10.00, and spline knot-spacing of 10.00 mm (Andersson et al., 2018). Final interpolation was performed using the jac method. The dMRI data was resampled to ACPC, generating an upsampled preprocessed dMRI data in ACPC space with 1.25 mm isotropic voxels.

Many internal operations of QSIPrep use Nilearn 0.10.1 (RRID:SCR_001362; Abraham et al., 2014) and Dipy (Garyfallidis et al., 2014). For more details of the pipeline, please see the section corresponding to workflows in QSIPrep's documentation.

#### Resting-state functional MRI preprocessing

Results included in this manuscript come from preprocessing performed using fMRIPrep 25.0.0 (RRID: SCR_016216; Esteban et al., 2019; Markiewicz et al., 2025), which is based on Nipype 1.9.2 (RRID: SCR_002502; Esteban et al., 2025; Gorgolewski et al., 2011).

A B0 nonuniformity map (or fieldmap) was estimated from the phase-drift maps measured with two consecutive GRE (gradient-recalled echo) acquisitions. The corresponding phase-map was phase-unwrapped with *prelude* (FSL 6.0.7.7; Jenkinson et al., 2012).

A total of one T1-weighted (T1w) image was found within the input BIDS dataset. The T1w image was corrected for intensity non-uniformity (INU) with N4BiasFieldCorrection (Tustison et al., 2010), distributed with ANTs 2.5.4 (RRID: SCR_004757; Avants et al., 2008) and used as T1w-reference throughout the workflow. The T1w-reference was then skull-stripped with a Nipype implementation of the antsBrainExtraction.sh workflow (from ANTs), using OASIS30ANTs as target template. Brain tissue segmentation of cerebrospinal fluid (CSF), white-matter (WM) and gray-matter (GM) was performed on the brain-extracted T1w using fast (FSL 6.0.7.7, RRID: SCR_002823; Zhang et al., 2001) Brain surfaces were reconstructed using FastSurfer 2.3.0 (Henschel et al., 2022, 2020), and the brain mask estimated previously was refined with a custom variation of the method to reconcile ANTs-derived and FastSurfer-derived segmentations of the cortical gray matter of Mindboggle (RRID:SCR_002438; Klein et al., 2017). Volume-based spatial normalization to MNI152NLin2009cAsym standard space was performed through nonlinear registration with antsRegistration (ANTs 2.5.4), using brain-extracted versions of both T1w reference and the T1w template. The following template was selected for spatial normalization and accessed with TemplateFlow (24.2.2; Ciric et al., 2022): ICBM 152 Nonlinear Asymmetrical template version 2009c [RRID: SCR_008796; TemplateFlow ID: MNI152NLin2009cAsym; Fonov et al., 2009]

For each of the one BOLD runs found per subject (across all tasks and sessions), the following preprocessing was performed. First, a reference volume was generated, using a custom methodology of fMRIPrep, for use in head motion correction. Head-motion parameters with respect to the BOLD reference (transformation matrices, and six corresponding rotation and translation parameters) are estimated before any spatiotemporal filtering using *mcflirt* (FSL; Jenkinson et al., 2002). FD and DVARS are calculated for each functional run, both using their implementations in Nipype (following the definitions by Power et al. 2014). The three global signals are extracted within the CSF, the WM, and the whole-brain masks. Additionally, a set of physiological regressors were extracted to allow for component-based noise correction (CompCor; Behzadi et al., 2007). Principal components are estimated after high-pass filtering the preprocessed BOLD time series (using a discrete cosine filter with 128s cut-off) for the two CompCor variants: temporal (tCompCor) and anatomical (aCompCor). tCompCor components are then calculated from the top 2% variable voxels within the brain mask. For aCompCor, three probabilistic masks (CSF, WM and combined CSF+WM) are generated in anatomical space. The implementation differs from that of Behzadi et al. in that instead of eroding the masks by two pixels on BOLD space, a mask of pixels that likely contain a volume fraction of GM is subtracted from the aCompCor masks. This mask is obtained by dilating a GM mask extracted from the FastSurfer's aseg segmentation, and it ensures components are not extracted from voxels containing a minimal fraction of GM. Finally, these masks are resampled into BOLD space and binarized by thresholding at 0.99 (as in the original implementation). Components are also calculated separately within the WM and CSF masks. For each CompCor decomposition, the *k* components with the largest singular values are retained, such that the retained components' time series are sufficient to explain 50 percent of variance across the nuisance mask (CSF, WM, combined, or temporal). The remaining components are dropped from consideration. The head-motion estimates calculated in the correction step were also placed within the corresponding confounds file. The confound time series derived from head motion estimates and global signals were expanded with the inclusion of temporal derivatives and quadratic terms for each (Satterthwaite et al., 2013). Frames that exceeded a threshold of 0.5 mm FD or 1.5 standardized DVARS were annotated as motion outliers. Additional nuisance timeseries are calculated by means of principal components analysis of the signal found within a thin band (crown) of voxels around the edge of the brain, as proposed by Patriat, Reynolds, and Birn (Patriat et al., 2017). All resamplings can be performed with a single interpolation step by composing all the pertinent transformations (i.e., head-motion transform matrices, susceptibility distortion correction, and co-registrations to

anatomical and output space). Gridded (volumetric) resamplings were performed using nitransforms, configured with cubic B-spline interpolation.
Many internal operations of fMRIPrep use Nilearn 0.11.1 (RRID: SCR_001362; Abraham et al., 2014) mostly within the functional processing workflow. For more details of the pipeline, see the section corresponding to workflows in fMRIPrep's documentation.

#### Resting-state functional MRI post-processing

The eXtensible Connectivity Pipeline- DCAN (XCP-D; Ciric et al., 2018; Mehta et al., 2024; Satterthwaite et al., 2013) version 0.10.7 was used to post-process the outputs of *fMRIPrep* version 25.0.0 (RRID: SCR_016216; Esteban et al., 2019; Markiewicz et al., 2025). XCP-D was built with *Nipype* version 1.10.0 (RRID: SCR_002502; Gorgolewski et al., 2011).

Native-space T1w images were transformed to MNI152NLin2009cAsym space at 1 mm3 resolution. For each of the one BOLD runs found per subject (across all tasks and sessions), the following post-processing was performed: FD was calculated from the motion parameters using the formula from Power et al. (2014), with automated head radius values based on subject brain masks. Volumes with framewise displacement greater than 0.5 mm were flagged as high-motion outliers for the sake of later censoring (Power et al., 2014). Nuisance regressors were selected according to the *'acompcor'* strategy. The top 5 aCompCor principal components from the white matter and cerebrospinal fluid compartments were selected as nuisance regressors (Behzadi et al., 2007), along with the six motion parameters and their temporal derivatives (Ciric et al., 2018; Satterthwaite et al., 2013). As the aCompCor regressors were generated on high-pass filtered data, the associated cosine basis regressors were included. This has the effect of high-pass filtering the data as well.

Nuisance regressors were regressed from the BOLD data using a denoising method based on *Nilearn*'s approach. Any volumes censored earlier in the workflow were first cubic spline interpolated in the BOLD data. Outlier volumes at the beginning or end of the time series were replaced with the closest low-motion volume's values, as cubic spline interpolation can produce extreme extrapolations. The time series were band-pass filtered using a second-order Butterworth filter, in order to retain signals between 0.008-0.1 Hz. The same filter was applied to the confounds. The resulting time series were then denoised via linear regression, in which the low-motion volumes from the BOLD time series and confounds were used to calculate parameter estimates, and then the interpolated time series were denoised using the low-motion parameter estimates. The interpolated time series were then censored using the temporal mask.

Many internal operations of *XCP-D* use *AFNI* (Cox, 1996; Cox and Hyde, 1997), *ANTS* (Avants et al., 2008), *TemplateFlow* version 24.2.2 (Ciric et al., 2022), *matplotlib* version 3.10.0 (Hunter, 2007), *Nibabel* version 5.3.2 (Brett et al., 2023), *Nilearn* version 0.11.1 (Abraham et al., 2014), *numpy* version 2.2.1 (Harris et al., 2020), *pybids* version 0.18.1 (Yarkoni et al., 2019), and *scipy* ve rsion 1.15.1 (Virtanen et al., 2020). For more details, see the *XCP-D* website (https://xcp-d.readthedocs.io).

**Supplementary Table 1. Mediation analyses testing whether individual TAP-M reaction time measures mediate the association between LC-nBM structural connectivity and cognitive dual-task cost.** Mediation analyses tested whether individual TAP-M reaction time measures explained the association between LC-nBM structural connectivity and cognitive dual-task cost. None of the indirect effects were significant after correction for multiple comparisons, whereas the direct and total effects remained significant. Thus, individual TAP-M measures did not mediate the association between LC-nBM connectivity and dual-task cost.

| Mediator | a path β, p | b path β, p | Direct effect c′, p | ACME / indirect effect, 95% CI, p | ADE, 95% CI, FDR-p | Total effect, 95% CI, FDR-p |
|---|---|---|---|---|---|---|
| Distractibility RT | 0.009, p = 0.263 | -0.028, p = 0.529 | 0.006, p = 0.021 | -0.0002 [-0.0021, 0.0007], p = 0.629 | 0.0059 [0.0022, 0.0108], p-FDR = 0.003 | 0.0057 [0.0023, 0.0100], p-FDR = 0.002 |
| Executive control RT | 0.025, p = 0.054 | -0.055, p = 0.050 | 0.007, p = 0.007 | -0.0014 [-0.0041, 0.0002], p = 0.121 | 0.0070 [0.0033, 0.0121], p-FDR = 0.001 | 0.0057 [0.0023, 0.0100], p-FDR = 0.002 |
| Divided attention visual RT | 0.033, p = 0.015 | 0.018, p = 0.512 | 0.005, p = 0.059 | 0.0006 [-0.0008, 0.0023], p = 0.397 | 0.0051 [0.0011, 0.0099], p-FDR = 0.016 | 0.0057 [0.0022, 0.0101], p-FDR = 0.002 |
| Divided attention auditory RT | 0.021, p = 0.111 | -0.021, p = 0.441 | 0.006, p = 0.019 | -0.0004 [-0.0023, 0.0007], p = 0.479 | 0.0061 [0.0023, 0.0106], p-FDR = 0.003 | 0.0057 [0.0022, 0.0100], p-FDR = 0.002 |
| Go correct RT | 0.009, p = 0.271 | -0.101, p = 0.019 | 0.007, p = 0.008 | -0.0009 [-0.0033, 0.0012], p = 0.376 | 0.0066 [0.0028, 0.0112], p-FDR = 0.002 | 0.0057 [0.0023, 0.0102], p-FDR = 0.002 |

Brett, M.A., Markiewicz, C.J.A., Hanke, M.A., Côté, M.-A.A., Cipollini, B.A., McCarthy, P.A., Jarecka, D.A., Cheng, C.P.A., Halchenko, Y.O.A., Cottaar, M.A., Larson, E.A., Ghosh, S.A., Wassermann, D.A., Gerhard, S.A., Lee, G.R.A., Wang, H.-T.A., Kastman, E.A., Kaczmarzyk, J.A., Guidotti, R.A., Daniel, J.A., Duek, O.A., Rokem, A.A., Madison, C.A., Papadopoulos Orfanos, D.A., Sólon, A.A., Moloney, B.A., Morency, F.C.A., Goncalves, M.A., Baratz, Z.A., Markello, R.A., Riddell, C.A., Burns, C.A., Millman, J.A., Gramfort, A.A., Leppäkangas, J.A., van den Bosch, J.J.F.A., Vincent, R.D.A., Braun, H.A., Subramaniam, K.A., Van, A.A., Gorgolewski, K.J.A., Raamana, P.R.A., Klug, J.A., Nichols, B.N.A., Baker, E.M.A., Hayashi, S.A., Pinsard, B.A., Haselgrove, C.A., Hymers, M.A., Esteban, O.A., Koudoro, S.A., Pérez-García, F.A., Dockès, J.A., Oosterhof, N.N.A., Amirbekian, B.A., Nimmo-Smith, I.A., Nguyen, L.A., Reddigari, S.A., St-Jean, S.A., Panfilov, E.A., Garyfallidis, E.A., Varoquaux, G.A., Legarreta, J.H.A., Hahn, K.S.A., Waller, L.A., Hinds, O.P.A., Fauber, B.A., Roberts, J.A., Poline, J.-B.A., Stutters, J.A., Jordan, K.A., Cieslak, M.A., Moreno, M.E.A., Hrnčiar, T.A., Haenel, V.A., Schwartz, Y.A., Darwin, B.C.A., Thirion, B.A., Gauthier, C.A., Solovey, I.A., Gonzalez, I.A., Palasubramaniam, J.A., Lecher, J.A., Leinweber, K.A., Raktivan, K.A., Calábková, M.A., Fischer, P.A., Gervais, P.A., Gadde, S.A., Ballinger, T.A., Roos, T.A., Reddam, V.R.A., freec84, 2023. nipy/nibabel: 5.0.1. Zenodo. https://doi.org/10.5281/zenodo.7633628

Ciric, R., Rosen, A.F.G., Erus, G., Cieslak, M., Adebimpe, A., Cook, P.A., Bassett, D.S., Davatzikos, C., Wolf, D.H., Satterthwaite, T.D., 2018. Mitigating head motion artifact in functional connectivity MRI. Nat. Protoc. 13, 2801–2826.

Ciric, R., Thompson, W.H., Lorenz, R., Goncalves, M., MacNicol, E.E., Markiewicz, C.J., Halchenko, Y.O., Ghosh, S.S., Gorgolewski, K.J., Poldrack, R.A., Esteban, O., 2022. TemplateFlow: FAIR-sharing of multi-scale, multi-species brain models. Nat. Methods 19,

1568–1571.
Cox, R.W., 1996. AFNI: software for analysis and visualization of functional magnetic resonance neuroimages. Comput. Biomed. Res. 29, 162–173.
Cox, R.W., Hyde, J.S., 1997. Software tools for analysis and visualization of fMRI data. NMR Biomed. 10, 171–178.
Esteban, O., Markiewicz, C.J., Blair, R.W., Moodie, C.A., Isik, A.I., Erramuzpe, A., Kent, J.D., Goncalves, M., DuPre, E., Snyder, M., Oya, H., Ghosh, S.S., Wright, J., Durnez, J., Poldrack, R.A., Gorgolewski, K.J., 2019. fMRIPrep: a robust preprocessing pipeline for functional MRI. Nat. Methods 16, 111–116.
Esteban, O., Markiewicz, C.J., Burns, C., Goncalves, M., Jarecka, D., Ziegler, E., Berleant, S., Ellis, D.G., Pinsard, B., Madison, C., Waskom, M., Notter, M.P., Clark, Daniel, Manhães-Savio, A., Halchenko, Y.O., Clark, Dav, Jordan, K., Dayan, M., Norgaard, M., Loney, F., Papadopoulos Orfanos, D., Salo, T., Johnson, H., Dewey, B.E., Bougacha, S., Keshavan, A., Yvernault, B., Christian, H., Hamalainen, C., Ćirić, R., Dubois, M., Joseph, M., Cipollini, B., Tilley, S., II, Visconti di Oleggio Castello, M., De La Vega, A., Wong, J., Kaczmarzyk, J., Huntenburg, J.M., Clark, M.G., Kent, J.D., Benderoff, E., Erickson, D., Dias, M. de F., Moloney, B., Hanke, M., Giavasis, S., Nichols, B.N., Tungaraza, R., Dell'Orco, A., Frohlich, C., Wassermann, D., de Hollander, G., Kepler, G., Koudoro, S., Eshaghi, A., Millman, J., Mancini, M., Close, T., Nielson, D.M., Varoquaux, G., Waller, L., Watanabe, A., Mordom, D., Cluce, J., Guillon, J., Robert-Fitzgerald, T., Chetverikov, A., Rokem, A., Acland, B., Bernardoni, F., Forbes, J., Markello, R., Gillman, A., Kong, X.-Z., Geisler, D., Salvatore, J., Espana, L., Gramfort, A., Doll, A., Buchanan, C., DuPre, E., Liu, S., Schaefer, A., Kleesiek, J., Butry, L., Sikka, S., Schwartz, Y., Ghayoor, A., Vaillant, G., Lee, J.A., Mattfeld, A., Richie-Halford, A., Liem, F., Perez-Guevara, M.F., Heinsfeld, A.S., Haselgrove, C., Durnez, J., Lampe, L., Poldrack, R., Glatard, T., Baratz, Z., Tabas, A., Cumba, C., Pérez-García, F., Genovese, M., Blair, R., Iqbal, S., Welch, D., Condamine, E., Contier, O., Triplett, W., Craddock, R.C., Correa, C., Stadler, J., Warner, J., Sisk, L.M., Falkiewicz, M., Sharp, P., Rothmei, S., Kim, S., Weinstein, A., Kahn, A.E., Kastman, E., Bottenhorn, K., Grignard, M., Perkins, L.N., Anijärv, T.E., Zhou, D., Bielievtsov, D., Cooper, G., Stojic, H., Hui Qian, T., Linkersdörfer, J., Renfro, M., Hinds, O., Stanley, O., Velasco, P., Küttner, R., Pauli, W.M., Wu, J., Xie, X., Glen, D., Kimbler, A., Meyers, B., Tarbert, C., Ginsburg, D., Haehn, D., Margulies, D.S., Ma, F., Malone, I.B., Snoek, L., Brett, M., Cieslak, M., Hallquist, M., Molina-Romero, M., Bilgel, M., Lee, N., Kuntke, P., Jalan, R., Inati, S., Gerhard, S., Mathotaarachchi, S., Saase, V., Van, A., Petre, B., Steele, C.J., Ort, E., Lerma-Usabiaga, G., Schwabacher, I., Arias, J., Lai, J., Pellman, J., Huguet, J., Junhao, W.E.N., Leinweber, K., Chawla, K., Weninger, L., Modat, M., Bannert, M.M., Mukhometzianov, R., Harms, R., Andberg, S.K., Matsubara, K., González Orozco, A.A., Routier, A., Marina, A., Davison, A., Floren, A., Park, A., Frederick, B., Cheung, B., McDermottroe, C., McNamee, D., Shachnev, D., Vogel, D., Flandin, G., Jones, H., Gonzalez, I., Varada, J., Schlamp, K., Podranski, K., Huang, L., Noel, M., Pannetier, N., Numssen, O., Khanuja, R., Urchs, S., Shim, S., Nickson, T., Huang, L., Broderick, W., Tambini, A., Mihai, P.G., Gorgolewski, K.J., Ghosh, S., 2025. nipy/nipype: 1.9.1. Zenodo. https://doi.org/10.5281/ZENODO.15054184
Fonov, V.S., Evans, A.C., McKinstry, R.C., Almli, C.R., Collins, D.L., 2009. Unbiased nonlinear average age-appropriate brain templates from birth to adulthood. NeuroImage 47, S102.
Garyfallidis, E., Brett, M., Amirbekian, B., Rokem, A., van der Walt, S., Descoteaux, M., Nimmo-Smith, I., Dipy Contributors, 2014. Dipy, a library for the analysis of diffusion MRI data. Front. Neuroinform. 8, 8.
Gorgolewski, K., Burns, C., Madison, C., Clark, D., Halchenko, Y., Waskom, M., Ghosh, S., 2011. Nipype: A Flexible, Lightweight and Extensible Neuroimaging Data Processing Framework in Python. Front. Neuroinform. 5.
Harris, C.R., Millman, K.J., van der Walt, S.J., Gommers, R., Virtanen, P., Cournapeau, D., Wieser, E., Taylor, J., Berg, S., Smith, N.J., Kern, R., Picus, M., Hoyer, S., van Kerkwijk, M.H.,

Brett, M., Haldane, A., del Río, J.F., Wiebe, M., Peterson, P., Gérard-Marchant, P., Sheppard, K., Reddy, T., Weckesser, W., Abbasi, H., Gohlke, C., Oliphant, T.E., 2020. Array programming with NumPy. Nature 585, 357–362.
Henschel, L., Conjeti, S., Estrada, S., Diers, K., Fischl, B., Reuter, M., 2020. FastSurfer - A fast and accurate deep learning based neuroimaging pipeline. Neuroimage 219, 117012.
Henschel, L., Kügler, D., Reuter, M., 2022. FastSurferVINN: Building resolution-independence into deep learning segmentation methods-A solution for HighRes brain MRI. Neuroimage 251, 118933.
Hoopes, A., Mora, J.S., Dalca, A.V., Fischl, B., Hoffmann, M., 2022. SynthStrip: skull-stripping for any brain image. Neuroimage 260, 119474.
Hunter, J.D., 2007. Matplotlib: A 2D Graphics Environment. Comput. Sci. Eng. 9, 90–95.
Jenkinson, M., Bannister, P., Brady, M., Smith, S., 2002. Improved Optimization for the Robust and Accurate Linear Registration and Motion Correction of Brain Images. NeuroImage 17, 825–841.
Jenkinson, M., Beckmann, C.F., Behrens, T.E.J., Woolrich, M.W., Smith, S.M., 2012. FSL. Neuroimage 62, 782–790.
Kellner, E., Dhital, B., Kiselev, V.G., Reisert, M., 2016. Gibbs-ringing artifact removal based on local subvoxel-shifts. Magn. Reson. Med. 76, 1574–1581.
Klein, A., Ghosh, S.S., Bao, F.S., Giard, J., Häme, Y., Stavsky, E., Lee, N., Rossa, B., Reuter, M., Chaibub Neto, E., Keshavan, A., 2017. Mindboggling morphometry of human brains. PLoS Comput. Biol. 13, e1005350.
Markiewicz, C.J., Esteban, O., Goncalves, M., Provins, C., Salo, T., Kent, J.D., DuPre, E., Ciric, R., Pinsard, B., Blair, R.W., Poldrack, R.A., Gorgolewski, K.J., 2025. fMRIPrep: a robust preprocessing pipeline for functional MRI. Zenodo. https://doi.org/10.5281/ZENODO.15085876
Mehta, K., Salo, T., Madison, T.J., Adebimpe, A., Bassett, D.S., Bertolero, M., Cieslak, M., Covitz, S., Houghton, A., Keller, A.S., Lundquist, J.T., Luo, A., Miranda-Dominguez, O., Nelson, S.M., Shafiei, G., Shanmugan, S., Shinohara, R.T., Smyser, C.D., Sydnor, V.J., Weldon, K.B., Feczko, E., Fair, D.A., Satterthwaite, T.D., 2024. XCP-D: A robust pipeline for the post-processing of fMRI data. Imaging Neuroscience 2, 1–26.
Patriat, R., Reynolds, R.C., Birn, R.M., 2017. An improved model of motion-related signal changes in fMRI. Neuroimage 144, 74–82.
Power, J.D., Mitra, A., Laumann, T.O., Snyder, A.Z., Schlaggar, B.L., Petersen, S.E., 2014. Methods to detect, characterize, and remove motion artifact in resting state fMRI. NeuroImage 84, 320–341.
Satterthwaite, T.D., Elliott, M.A., Gerraty, R.T., Ruparel, K., Loughead, J., Calkins, M.E., Eickhoff, S.B., Hakonarson, H., Gur, R.C., Gur, R.E., Wolf, D.H., 2013. An improved framework for confound regression and filtering for control of motion artifact in the preprocessing of resting-state functional connectivity data. NeuroImage 64, 240–256.
Tournier, J.-D., Smith, R., Raffelt, D., Tabbara, R., Dhollander, T., Pietsch, M., Christiaens, D., Jeurissen, B., Yeh, C.-H., Connelly, A., 2019. MRtrix3: A fast, flexible and open software framework for medical image processing and visualisation. Neuroimage 202, 116137.
Tustison, N.J., Avants, B.B., Cook, P.A., Zheng, Y., Egan, A., Yushkevich, P.A., Gee, J.C., 2010. N4ITK: Improved N3 Bias Correction. IEEE Trans. Med. Imaging 29, 1310–1320.
Veraart, J., Novikov, D.S., Christiaens, D., Ades-Aron, B., Sijbers, J., Fieremans, E., 2016. Denoising of diffusion MRI using random matrix theory. Neuroimage 142, 394–406.
Virtanen, P., Gommers, R., Oliphant, T.E., Haberland, M., Reddy, T., Cournapeau, D., Burovski, E., Peterson, P., Weckesser, W., Bright, J., van der Walt, S.J., Brett, M., Wilson, J., Millman, K.J., Mayorov, N., Nelson, A.R.J., Jones, E., Kern, R., Larson, E., Carey, C.J., Polat, İ., Feng, Y., Moore, E.W., VanderPlas, J., Laxalde, D., Perktold, J., Cimrman, R., Henriksen, I., Quintero, E.A., Harris, C.R., Archibald, A.M., Ribeiro, A.H., Pedregosa, F., van Mulbregt, P., Vijaykumar, A., Bardelli, A.P., Rothberg, A., Hilboll, A., Kloeckner, A., Scopatz, A., Lee, A., Rokem, A.,

Woods, C.N., Fulton, C., Masson, C., Häggström, C., Fitzgerald, C., Nicholson, D.A., Hagen, D.R., Pasechnik, D.V., Olivetti, E., Martin, E., Wieser, E., Silva, F., Lenders, F., Wilhelm, F., Young, G., Price, G.A., Ingold, G.-L., Allen, G.E., Lee, G.R., Audren, H., Probst, I., Dietrich, J.P., Silterra, J., Webber, J.T., Slavič, J., Nothman, J., Buchner, J., Kulick, J., Schönberger, J.L., de Miranda Cardoso, J.V., Reimer, J., Harrington, J., Rodríguez, J.L.C., Nunez-Iglesias, J., Kuczynski, J., Tritz, K., Thoma, M., Newville, M., Kümmerer, M., Bolingbroke, M., Tartre, M., Pak, M., Smith, N.J., Nowaczyk, N., Shebanov, N., Pavlyk, O., Brodtkorb, P.A., Lee, P., McGibbon, R.T., Feldbauer, R., Lewis, S., Tygier, S., Sievert, S., Vigna, S., Peterson, S., More, S., Pudlik, T., Oshima, T., Pingel, T.J., Robitaille, T.P., Spura, T., Jones, T.R., Cera, T., Leslie, T., Zito, T., Krauss, T., Upadhyay, U., Halchenko, Y.O., Vázquez-Baeza, Y., SciPy, C., 2020. SciPy 1.0: fundamental algorithms for scientific computing in Python. Nat. Methods 17, 261–272.

Yarkoni, T., Markiewicz, C.J., de la Vega, A., Gorgolewski, K.J., Salo, T., Halchenko, Y.O., McNamara, Q., DeStasio, K., Poline, J.B., Petrov, D., Hayot-Sasson, V., Nielson, D.M., Carlin, J., Kiar, G., Whitaker, K., DuPre, E., Wagner, A., Tirrell, L.S., Jas, M., Hanke, M., Poldrack, R.A., Esteban, O., Appelhoff, S., Holdgraf, C., Staden, I., Thirion, B., Kleinschmidt, D.F., Lee, J.A., Visconti di Oleggio Castello, M., Notter, M.P., Blair, R., 2019. PyBIDS: Python tools for BIDS datasets. J. Open Source Softw. 4.

Zhang, Y., Brady, M., Smith, S., 2001. Segmentation of brain MR images through a hidden Markov random field model and the expectation-maximization algorithm. IEEE Trans. Med. Imaging 20, 45–57.
